# Frequency-dependent representation of reinforcement-related information in the human medial and lateral prefrontal cortex

**DOI:** 10.1101/029058

**Authors:** Elliot H. Smith, Garrett P. Banks, Charles B. Mikell, Sydney S. Cash, Shaun R. Patel, Emad A. Eskandar, Sameer A. Sheth

## Abstract

The feedback-related negativity (FRN) is a commonly observed potential in scalp electroencephalography (EEG) studies related to the valence of feedback about a subject’s performance. This potential classically manifests as a negative deflection in medial frontocentral EEG contacts following negative feedback. Recent work has shown prominence of theta power in the spectral composition of the FRN, placing it within the larger class of “frontal midline theta” cognitive control signals. Although the dorsal anterior cingulate cortex (dACC) is thought to be the cortical generator of the FRN, conclusive data regarding its origin and propagation are lacking. Here we examine intracranial electrophysiology from the human medial and lateral prefrontal cortex (PFC) in order to better understand the anatomical localization and communication patterns of the FRN. We show that the FRN is evident in both low- and high-frequency local field potentials (LFPs) recorded on electrocorticography. The FRN is larger in medial compared to lateral PFC, and coupling between theta band phase and high frequency LFP power is also greater in medial PFC. Using Granger causality and conditional mutual information analyses, we provide evidence that feedback-related information propagates from medial to lateral PFC, and that this information transfer oscillates with theta-range periodicity. These results provide evidence for the dACC as the cortical source of the FRN, provide insight into the local computation of frontal midline theta, and have implications for reinforcement learning models of cognitive control.

**Significance Statement**: Using intracranial electrophysiology in humans, this work addresses questions about a frequently studied feedback-related electroencephalographic signal, illuminating anatomical and functional properties of the representation of feedback-related reinforcement during decision-making across the medial to lateral extent of the human prefrontal cortex.

## Introduction

Modifying future behavior based on reinforcement is essential for survival in complex environments. A prominent event-related potential (ERP) observed on EEG is the feedback-related negativity (FRN), which manifests as the difference between ERPs evoked by positive and negative feedback (Walsh and Anderson, 2012). The FRN occurs approximately 250 ms after feedback, regardless of the feedback modality (Miltner et al., 1997).

The FRN is most commonly observed on frontocentral EEG contacts, where its amplitude is highest (Miltner et al., 1997; Luu et al., 2003; Silvetti et al., 2014), but has also been observed on parietal contacts (Cohen and Ranganath, 2007; Pfabigan et al., 2010). EEG and MEG source localization has commonly identified the dorsomedial prefrontal cortex (dmPFC), especially the dorsal anterior cingulate cortex (dACC), as the source of the FRN (Miltner et al., 2003; Herrmann et al., 2004; Luu et al., 2004; Roger et al., 2010; Doñamayor et al., 2011; Walsh and Anderson, 2013). fMRI studies have found areas that have increased BOLD signal for positive feedback, compared with negative feedback (Nieuwenhuis et al., 2005b). These studies have implicated a broad network in FRN generation, including cingulate cortex, dorsolateral PFC (dlPFC) the basal ganglia, and amygdalae (Holroyd et al., 2004b). One recent study that combined the temporal resolution of EEG with the spatial resolution of fMRI localized the FRN to either the dACC specifically, or to a distributed frontal “salience” network, depending on the analysis technique used (Hauser et al., 2013).

An influential account of FRN function based on reinforcement learning concepts links it to the reward system (Holroyd and Coles, 2002a; Holroyd et al., 2008). Mesencephalic dopamine projections to the striatum and dmPFC reinforce advantageous behaviors (Schultz, 2002). Phasic dopamine release is thought to encode reward prediction error (RPE) signals, or differences between expected and actual rewards (Schultz, 2013). Errors, resulting in negative feedback or reward omission, produce phasic decreases in dopamine release. As dopaminergic input is inhibitory to the dACC, phasic decreases disinhibit layer V dACC neurons, allowing them to depolarize and produce ERPs (Holroyd and Coles, 2002b). Indeed, FRN magnitude is directly proportional to RPE magnitude and the amount of behavioral adaptation in subsequent trials (Cavanagh et al., 2010).

Recent studies have shown that the FRN, along with other frontocentral negativities such as the error-related negativity (ERN) and N2, share common features. Spectral analyses have demonstrated increased 4-7 Hz power, suggesting a common mechanism of communication for these frontal midline theta (FMΘ) signals (Holroyd et al., 2002; Chase et al., 2011; Cavanagh and Frank, 2014; Cavanagh and Shackman, 2014). These signals are thought to convey aspects of cognitive control and may have evolved as reactions to threatening situations, engaging cognitive resources in times of high demand (Cavanagh and Shackman, 2014).

Here we examine intracranial electrophysiology from the human dACC and dlPFC to localize and characterize the neural representation of prefrontal feedback signals. We show that the dACC is the source of the FRN, that the information contained in this signal propagates to dlPFC, and that this communication occurs in the theta frequency band.

## Materials and Methods

*Ethics statement.* The Columbia University Medical Center and Massachusetts General Hospital Institutional Review Boards approved these experiments. All patients in this study provided informed consent prior to participating in this research.

*Subjects.* We examined intracranial electrocorticographic (ECoG) signals in 7 patients (2 female) undergoing monitoring for medically refractory epilepsy. Eight-contact depth electrodes (AdTech Medical, Racine, WI) were implanted through the mediolateral extent of prefrontal cortex (10 left hemisphere and 9 right hemisphere), thus providing simultaneous LFP recordings from dACC and dlPFC. Electrode placement was determined solely on clinical grounds, and participants were free to withdraw from the study without influencing their clinical care.

*Depth Electrode localization.* Depth electrodes were localized using pre-operative magnetic resonance images (MRI) and post-implant computed tomography (CT) scans. A linear transform from CT space to pre-operative MRI space was designed using FMRIB Software Library’s (FSL) linear transformation algorithm (Jenkinson and Smith, 2001). Next, a transform from patient MRI space to 2 mm Montreal Neurological Institute standard space using both linear and non-linear transforms was implemented (Jenkinson et al., 2005; Andersson et al., 2013). Depth electrodes’ medial and lateral-most contacts were then respectively superimposed on the medial and lateral surface of the standard brain for visualization.

*Behavioral task.* Recordings were acquired while patients performed the Multi-Source Interference Task (MSIT) (Figure 1A) (Sheth et al., 2012). The MSIT is a Stroop-like cognitive interference task requiring the subject to view a stimulus consisting of three numbers (composed of integers 0 through 3), and identify the one number (“target”) different from the other two (“distracters”). The subject has to indicate his/her choice on a 3-button pad. If the target number is “1”, the left button is the correct choice; if “2”, the middle button; and if “3”, the right button. Importantly, the subject must choose the correct button regardless of where the target number appears in the sequence. Reaction time (RT) was defined as the time between stimulus presentation and subject response. Differences in RT among conditions was tested for significance using a one way Kruskal-Wallis test over interference types, and a Wilcoxon signed rank test between RTs for feedback conditions.

**Figure 1.**
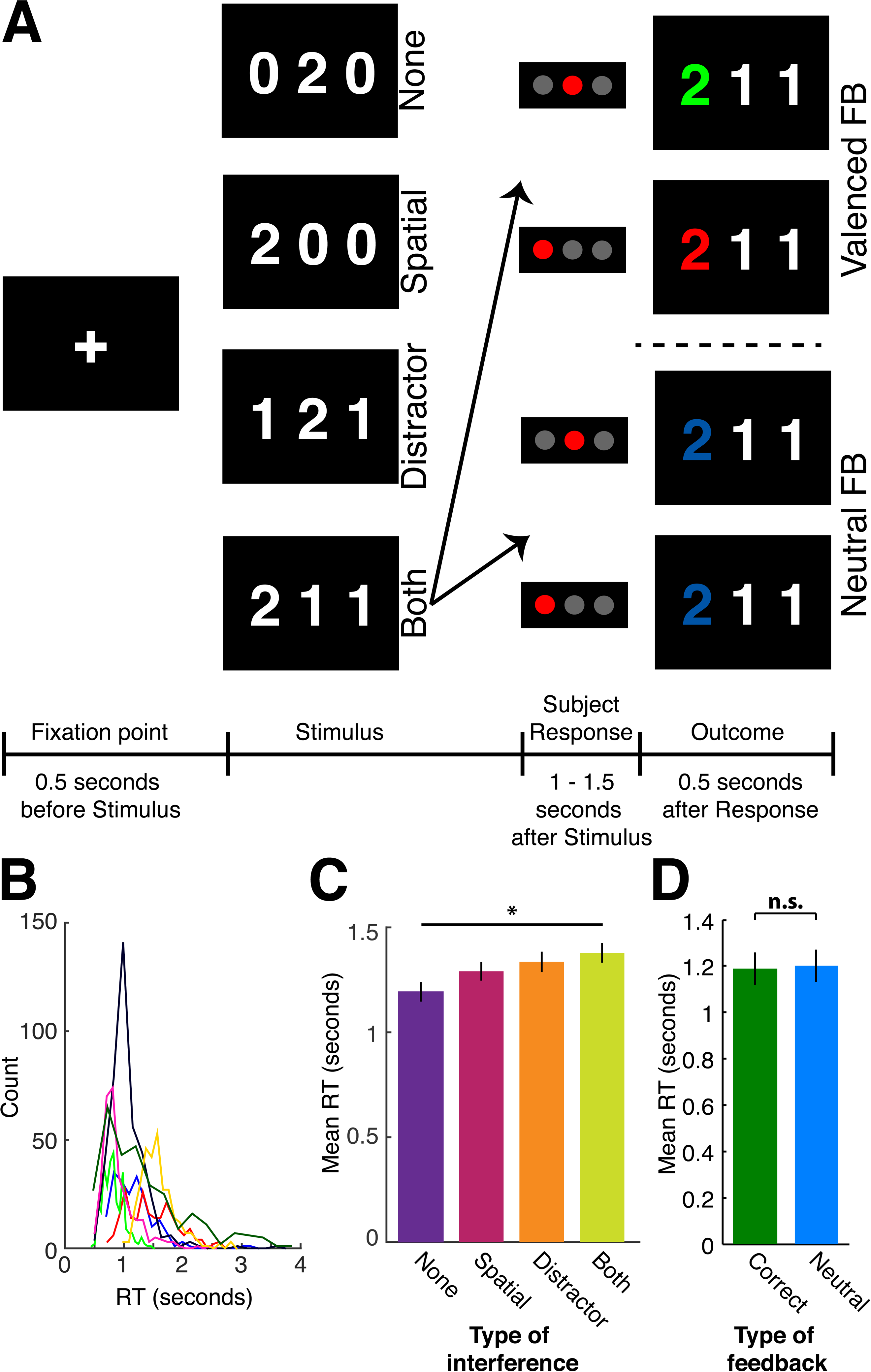
Task description and behavioral performance. **A,** MSIT task description and timeline. A fixation cross appears on the screen 500 milliseconds before stimuli are shown. Four labeled examples of the different types of interference are shown above the stimulus period in the task timeline. Feedback occurs following the subject’s response to the stimulus. In valenced feedback trials, the target number turns green for correct responses and red for incorrect responses. In neutral feedback trials, the target number turns blue regardless of whether the subject responded correctly. Examples of the four possible types of feedback are shown for the example stimulus that contains both types of interference. **B,** Distributions of RTs for each subject colored by subject. **C,** Mean RTs across subjects for trials containing the four different types of interference. The asterisk indicates statistical significance over conditions for the ANOVA. **D,** Mean RTs across subjects for trials following correct and neutral feedback. “n.s.” indicates no significant difference.

The task contains two types of cognitive interference. Simon interference occurs due to the presence of spatial incongruence between the position of the target number in the stimulus and the position of the correct button on the button pad. For example, the correct choice for both “200” and “020” is the middle button; the first cue contains spatial interference (incongruence between the left position of the target number in the stimulus and middle position of the correct button choice), whereas the second one does not. Eriksen Flanker interference occurs due to the presence of distracters that are possible button choices. For example, the correct choice for both “133” and “100” is the left button; the first cue contains distracter interference (“3” is a possible button choice), whereas the second one does not (“0” is not a possible choice).

Patients received either valenced or neutral feedback in blocks of 10 trials. During valenced feedback trials, the target number changes color to indicate whether the patient responded correctly: green for correct responses and red for incorrect responses. During neutral feedback trials, the target number changes to blue regardless of the response (Figure 1A).

*Data collection and preprocessing.* Neural data from depth electrodes were sampled at 500 Hz on the clinical data acquisition system (Xltek, Natus Medical Inc., San Carlos, CA). First, removal of the common mode was achieved by reconstructing the data without the first principal component, based on the covariance of all examined contacts (including white matter contacts) in each patient. This method removed DC offsets, common artifacts, and line noise present in the raw data. Spectrograms were then calculated using multi-tapered spectral analysis with a time-bandwidth product of 5 and 9 tapers, 200 ms windows and 10 ms step sizes, and normalized relative to mean baseline spectra (500 ms prior to the appearance of the fixation point). High gamma activity was derived from the mean values of spectrograms between 70 and 125 Hz. These frequency limits were chosen to be above the line noise on the low end, and half of the Nyquist frequency on the upper end. To generate low-frequency ERPs, data were low-pass filtered at 40 Hz (150^th^ order Fir filter) and averaged over trials.

In order to motivate our frequency-specific hypotheses and compare this study with previous EEG studies, spectrograms and inter-trial phase coherence (ITC) were calculated from Morlet wavelet decompositions on 72 scales from 1 to 125 Hz. The group spectrogram is the average of the absolute value of each subject’s spectrogram. ITC was derived from averaging the absolute value of the phase of the LFP at each frequency value in the wavelet decomposition over trials. Paired t-tests over trials were used to assess significant differences between feedback conditions during the outcome period for each frequency band in the time-frequency representations. These results were corrected for the number of frequency bands using the Benjamini-Hochberg false discovery rate (FDR) (Hochberg and Tamhane, 1987).

*FRN quantification.* The FRN for both low and high frequency signals was defined as the difference in signal between correct and neutral feedback trials. We did not calculate the difference between correct and incorrect feedback responses because of the low error rate (see below). Previous work has shown that the FRN is dependent upon expectations established by the context, such that an FRN is observed for negative feedback when the range of outcomes varies from neutral to negative, but for neutral feedback when the range varies from positive to neutral (Holroyd et al., 2004a; Nieuwenhuis et al., 2005a; Jessup et al., 2010).

To account for variation in the size of the mean ERP between medial and lateral sites, low frequency FRN amplitude was quantified by normalizing the maximum FRN amplitude by the ERP amplitude (i.e the difference between the minimum and maximum) at each contact. FRN amplitude is therefore reported relative to the size of the mean ERP at each site. For high frequency signals, the FRN was quantified by examining differences in high gamma power during feedback presentation. Hypothesis testing for these results were carried out using Mann-Whitney U tests within patients and between feedback conditions for both low frequency signals and high gamma power. In order to address causal influences of feedback-related representations in dACC and dlPFC, we calculated the latency of the FRN signal as the point of separation of the averaged high gamma traces for neutral and correct feedback as long as that separation was maintained for at least 500 ms. Latency was further quantified as the peak cross-correlation of evoked high gamma signals between medial and lateral contacts.

*Phase-amplitude coupling.* To determine whether there was a functional interaction between low and high frequency LFPs we examined phase-amplitude coupling between a range of low frequency (2-25 Hz) phases and high-gamma power for each channel separately. The high-gamma power was determined with the Hilbert transform of the band-pass filtered signal between 70 and 125 Hz (4^th^ order butterworth). Frequencies for phase were filtered with a 3 Hz-wide band-pass filter centered on each phase frequency (4^th^ order butterworth). Phase values were extracted using the arctangent of the real and imaginary values of the Hilbert transforms of each filtered low-frequency band. High gamma amplitudes were then binned in 32 phase bins between -! and ! for each low-frequency band and retained for further analysis. To determine significantly modulated phase frequencies, each phase frequency’s distribution of high gamma amplitudes was tested for significant modulation with a Kolmogorov-Smirnov goodness-of-fit test against a uniform null distribution and corrected for the number of frequency bands using the FDR (Hochberg and Tamhane, 1987). The best frequency-for-phase for each electrode and patient was also determined from the Kolmogorov-Smirnov goodness-of-fit test, as the most significant frequency-for-phase band.

In order to quantify the amount of modulation in significant frequency-for-phase bands, modulation indices were calculated for significantly modulated frequency-for-phase bands. Modulation index was calculated in a similar manner to Canolty et al. (Canolty et al., 2006) and Miller et al. (Miller et al., 2010; 2012). Briefly, the analytic signals returned from the Hilbert transforms of the filtered ECoG data for each trial, n, were separated into their amplitude and phase components *a_n_* and *ϕ_n_*. The modulation index used here is the magnitude of the vector defined by the complex variable

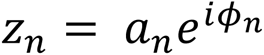

during the feedback period. Significance was determined by comparing distributions of these vectors projected onto the mean vector for each feedback condition using ANOVA, and the Tukey-Kramer test for pairwise comparisons.

*Granger Causality.* Granger causality (GC) is a commonly used metric to infer directional connectivity among brain areas (Seth, 2010); (Boatman-Reich et al., 2010); (Barnett and Seth, 2014; Rodrigues and Andrade, 2014). GC measures the extent to which one signal can predict another by imposing time lags on one signal and examining resulting changes in the regression coefficients of an autoregressive model of both (Granger, 1969; Seth and Edelman, 2007). Here within subject GC was determined using the Multivariate Granger Causality Toolbox for Matlab (Barnett and Seth, 2014). We examined all possible interactions between the two medial- and lateral-most contacts among all electrodes in each patient, for the feedback period. We considered interactions between combinations of pairs of electrodes in this frontal network (630 possible connections). The Lyapunov exponents of the autoregressive model based on the raw data were initially greater than one, indicating that the data did not meet the criteria of stationarity required for GC analysis. A μ-law compressor, with μ = 255, was therefore applied to the z-scored broadband time series to compress the dynamic range of the signals while maintaining the phase structure necessary to infer directional connectivity. The multivariate autoregressive model, *A*, henceforth had a stable spectral radius (*ρ*(*A*) = 0.96). Appropriate model order specification was determined using the Bayesian information criterion (Bressler and Seth, 2011). A model order of 22 was determined to be optimal for each of the seven patients and the mean± standard deviation (over the seven patients) number of lags used was 104.34±36.71 samples.

Statistical significance was determined by testing the distribution generated by the multivariate autoregressive model against a theoretical *χ*^2^-distribution. The criterion for significance was set at 0.05 after correction for multiple hypotheses (Hochberg and Tamhane, 1987).

*Permutation Conditional Mutual Information.* GC analysis relies on several assumptions that may not always hold in ECoG data, including stationarity, as mentioned above. Conditional mutual information (CMI) was therefore examined as an additional measure of information transfer between medial and lateral contacts. CMI is an information theoretic measure of information transfer between cortical areas or recording sites. CMI has been used to describe information transfer in cortical circuits without relying on Gaussian statistics or imposing time delays on neural data (Quinn et al., 2010; Salvador et al., 2010; Smith et al., 2012; 2013).

Here we computed the permutation, or symbolic, CMI (Li and Ouyang, 2010; King et al., 2013). For permutation CMI, broadband time series were discretized into groups of 6 symbols, 3 samples at a time, representing changes in the recorded voltage on each contact. Probability distributions were then generated from the pairwise frequencies of symbols derived from the two signals being compared (Keller and Lauffer, 2003; Zunino et al., 2010). CMI was then computed as

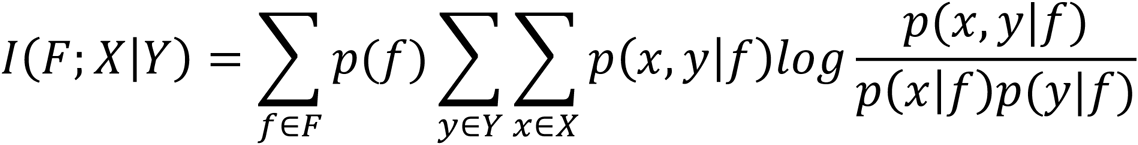

Where F represents feedback, *X* is the response in one cortical area and *Y* is the response in another cortical area. *p(f)* is the probability a trial contains feedback*; p(x)* and *p(y)* are the probabilities of observing responses *x* and *y* respectively.

We calculated CMI for epochs during and just before the feedback period for medial- and lateral-most pairs of contacts on each electrode. Symbolic CMI therefore tells us how much information a signal contains about feedback valence, given that we know the response to feedback valences on the other end of the electrode. CMI regarding feedback valence was calculated for the medial electrode given that the response in the lateral electrode was known and then for the lateral electrode given that the response in the medial electrode was known. This calculation was made in 250 ms windows sliding 5 ms in time for 4 seconds surrounding feedback presentation.

We also calculated CMI for the same data with randomly shuffled samples for use in statistical comparisons. Permutation CMI is a relative measure of information transfer. Statistically significant differences from shuffled data were therefore interpreted as representing information transfer from one cortical area to the other cortical area. Statistical significance was assessed within patients using a modified *χ*^2^ test, comparing *χ*^2^ distributions generated from 2*Nlog*_2_ times the lateral-to-medial and medial-to-lateral CMI to the *χ*^2^ distribution from the shuffled data (Fan et al., 2000). Here *N* is the number of trials, and the degrees of freedom for each distribution are based on the number of possible responses for each stimulus presentation and the total number of presentations (Ince et al., 2011).

To quantify the amplitude and phase of oscillations observed in the CMI signal, multi-taper spectra (5 leading tapers and a time-bandwidth product of 3) of the CMI signals were calculated.

## Results

### Behavioral data

Seven subjects performed the task with similar behavior as has been previously reported, with RT increasing with amount of interference in the stimulus (Figures 1B and C) (Horga et al., 2011; Sheth et al., 2012). There was no significant difference in RT between trials with valenced vs. neutral feedback (Wilcoxon signed rank, P = 0.78, N = 7) (Figure 1D). Subjects had a 1.6± 1.3 % error rate over an average of 289±65.3 trials per subject.

### Depth electrode localization

Each depth electrode had 8 contacts. In most electrodes, the two medial- and lateral-most contacts were determined to be within gray matter. The middle four contacts in each electrode were predominantly in white matter and therefore were not examined. Two medial contacts on one electrode were also determined to be in white matter. This electrode was excluded from further analysis. We therefore examined responses on 72 cortical contacts over all 7 patients. Figure 2 shows the medial and lateral most contacts in each patient projected onto the medial and lateral surfaces of the MNI standard brain.

**Figure 2.**
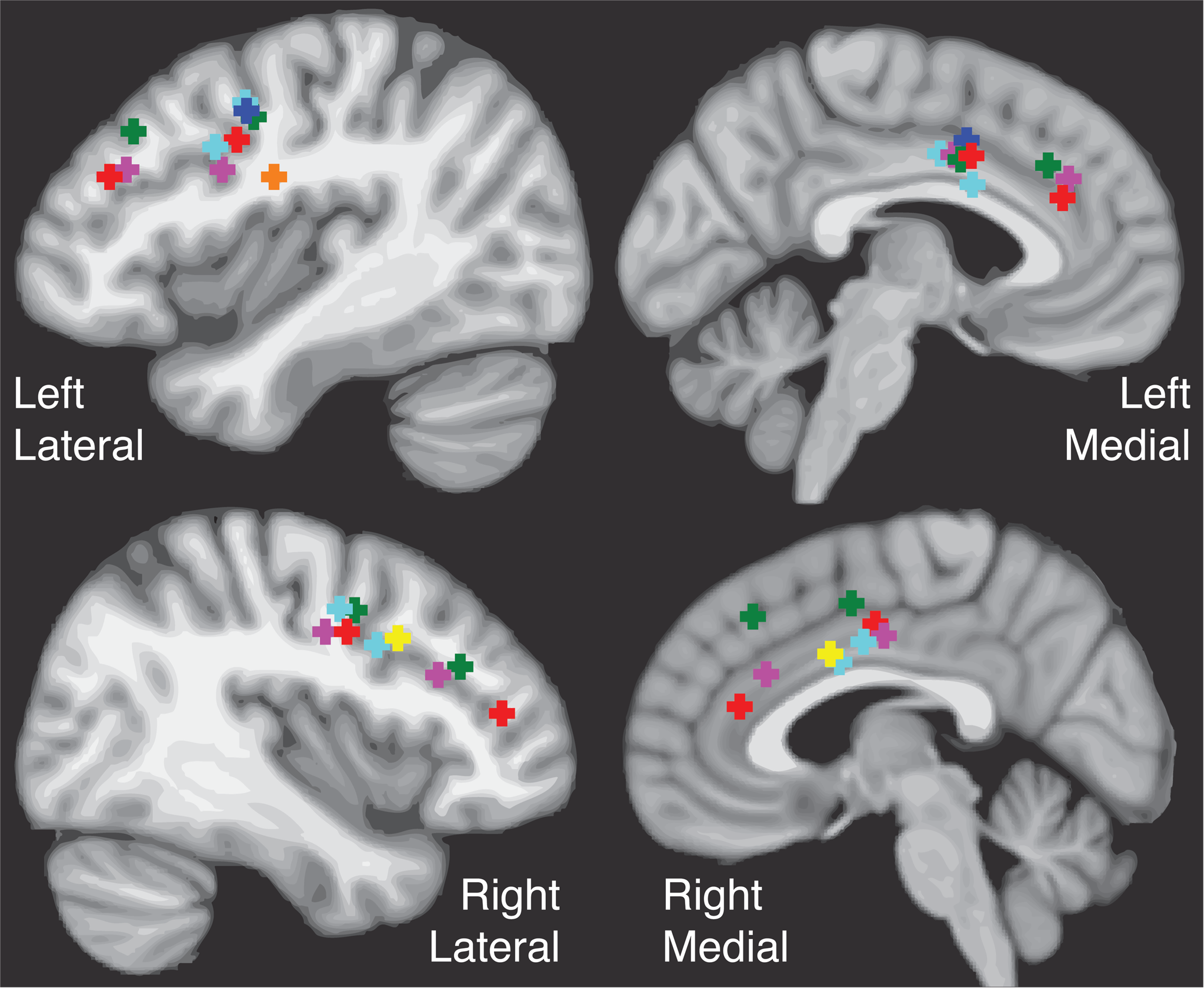
Depth Electrode locations in seven subjects. The locations of the medial and lateral–most contacts on each depth electrode projected onto the medial and lateral surfaces of each hemisphere of the MNI standard brain. Colors represent individual subjects.

### FRN is evident in high and low frequency signals and is larger in dACC than dlPFC

We found evidence for differential representations of feedback valence in both low and high frequency components of the ECoG recordings (Figures 3A and 3B; Paired t-tests, all p < 0.01). Figure 3C depicts a representative feedback-locked ERP demonstrating the low-frequency FRN. This intracranial FRN was significant in the low frequency averaged LFP across subjects (Kruskal-Wallis one-way ANOVA, p << 0.01, df = 71 contacts). There was a main effect of feedback valence in 32 of 36 medial electrodes and 32 of 36 lateral electrodes (Tukey-Kramer, all p = 0.0013, N = 36 contacts each), indicating that the FRN commonly observed on scalp EEG is evident in intracranial recordings. Low frequency FRN was significantly greater in medial (dACC) contacts than lateral (dlPFC) contacts (Mann-Whitney U test, p = 0.0041, N = 32 contacts) (Figure 3E).

**Figure 3.**
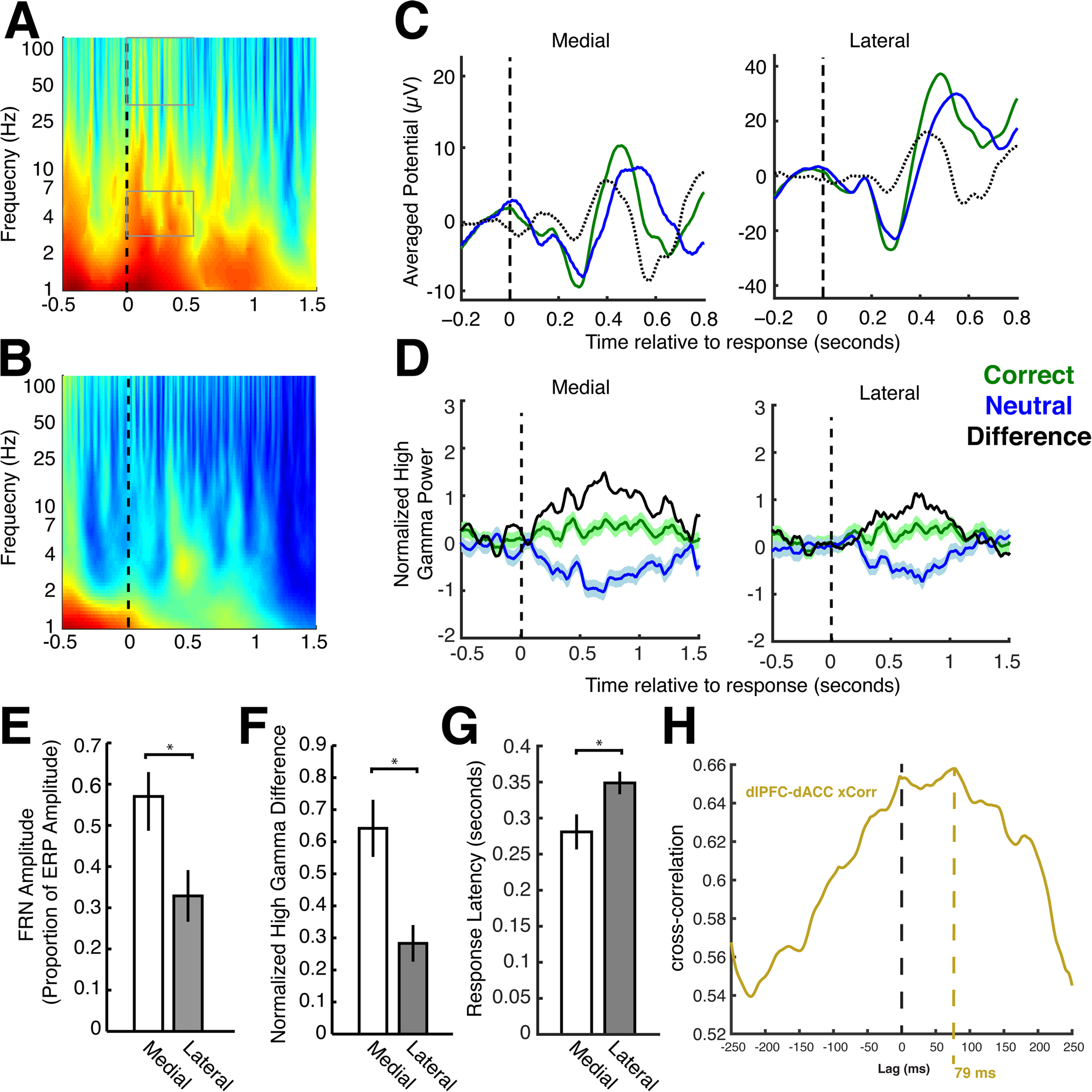
FRN in high and low frequency LFP. **A,** Response aligned spectrograms averaged over trials (mean±S.D. = 289±65.32) and patients. Frequency bands during the outcome period that are significantly different between feedback conditions arte outlined with gray boxes (paired t-tests, corrected for 72 hypotheses, all p<0.01). **B,** Response aligned inter-trial phase coherence averaged over trials and patients. **C,** Representative event related potentials from one subject for medial and lateral electrode contacts. Green and blue lines represent correct and neutral feedback conditions, respectively, and black lines represent the difference. Shading represents standard errors. **D,** Representative evoked high gamma response from one subject. **E,** Quantification of FRN amplitude from ERP (Mann-Whitney U test, p = 0.0041, N = 32 contacts). **F,** Quantification of high gamma power for each feedback condition (Mann-Whitney U test, p = 0.00096, N = 22 contacts). **G,** Quantification of latency differences between medial and lateral high gamma responses (two-sample t-test, p = 0.02, N = 22 contacts). Asterisks indicate significant differences (p<0.05). **H,** Lagged cross-correlogram between medial and lateral high gamma responses. The peak indicated that mean high gamma responses in lateral contacts lagged those in medial contacts by 79 ms.

Given the relationship between high-gamma ECoG signals and local neuronal processing (Manning et al., 2009; Miller et al., 2009; Ray and Maunsell, 2011; Buzsáki et al., 2012; Einevoll et al., 2013), we also examined the high-gamma representation of the FRN in dACC and dlPFC (Figure 3D). Feedback significantly affected high-gamma responses across subjects (Kruskal-Wallis one-way ANOVA, p << 0.01, df = 71 contacts). Within subjects, and across trials, there were also significant differences in evoked high-gamma power between correct and neutral feedback trials in 34 of 36 medial electrodes and 22 of 36 lateral contacts. As with low frequency potentials, a medial-to-lateral decreasing gradient in FRN amplitude was observed in high-gamma responses (Mann-Whitney U test, p = 0.00096, N = 22 contacts) (Figure 3F). Furthermore, latency of the FRN response was shorter in medial compared to lateral contacts (two-sample t-test, p = 0.02, N = 22 contacts) (Figure 3G). Cross-correlation of medial and lateral high gamma signals corroborated this result by estimating the mean latency as 79 ms over patients (Figure 3H). These results indicate that the FRN is likely generated medially, in the dACC.

### High and low frequency LFPs are functionally related, more so in medial PFC

We next examined coupling between low frequency (2 – 25 Hz) LFP phase and high frequency (70 – 125 Hz) LFP amplitude. We found significant phase-amplitude coupling (PAC) between theta phase and high gamma amplitude in all subjects (Kolmogorov-Smirnov goodness-of-fit test, all p < 0.01, N = 50 contacts each, corrected for FDR due to 23 null hypotheses) (Figure 4A, B). The peak frequency-for-phase was 5.1 ±0.2 Hz for correct feedback trials and 4.8±0.4 Hz for neutral feedback. There was no significant difference in best frequency-for-phase between medial and lateral contacts (Kruskal-Wallis one-way ANOVA, p = 0.27, dF = 71 contacts). The magnitude of the theta-high gamma coupling, as measured by the modulation index, was significantly greater for correct feedback trials than for neutral feedback trials in medial (Tukey-Kramer, p = 0.0052, N = 36 contacts), but not in lateral (Tukey-Kramer, p = 0.0598, N = 36 contacts) contacts (Figure 4C). In addition, the difference in theta-high gamma coupling between correct and neutral trials was greater in medial contacts than lateral contacts (Tukey-Kramer, p = 0.0098, N = 36 contacts) (Figure 4E). These results tie the local activity of the dACC and PFC to the theta rhythm that can be observed on scalp EEG and show that feedback information is also represented in the strength of theta-gamma coupling in dACC.

**Figure 4.**
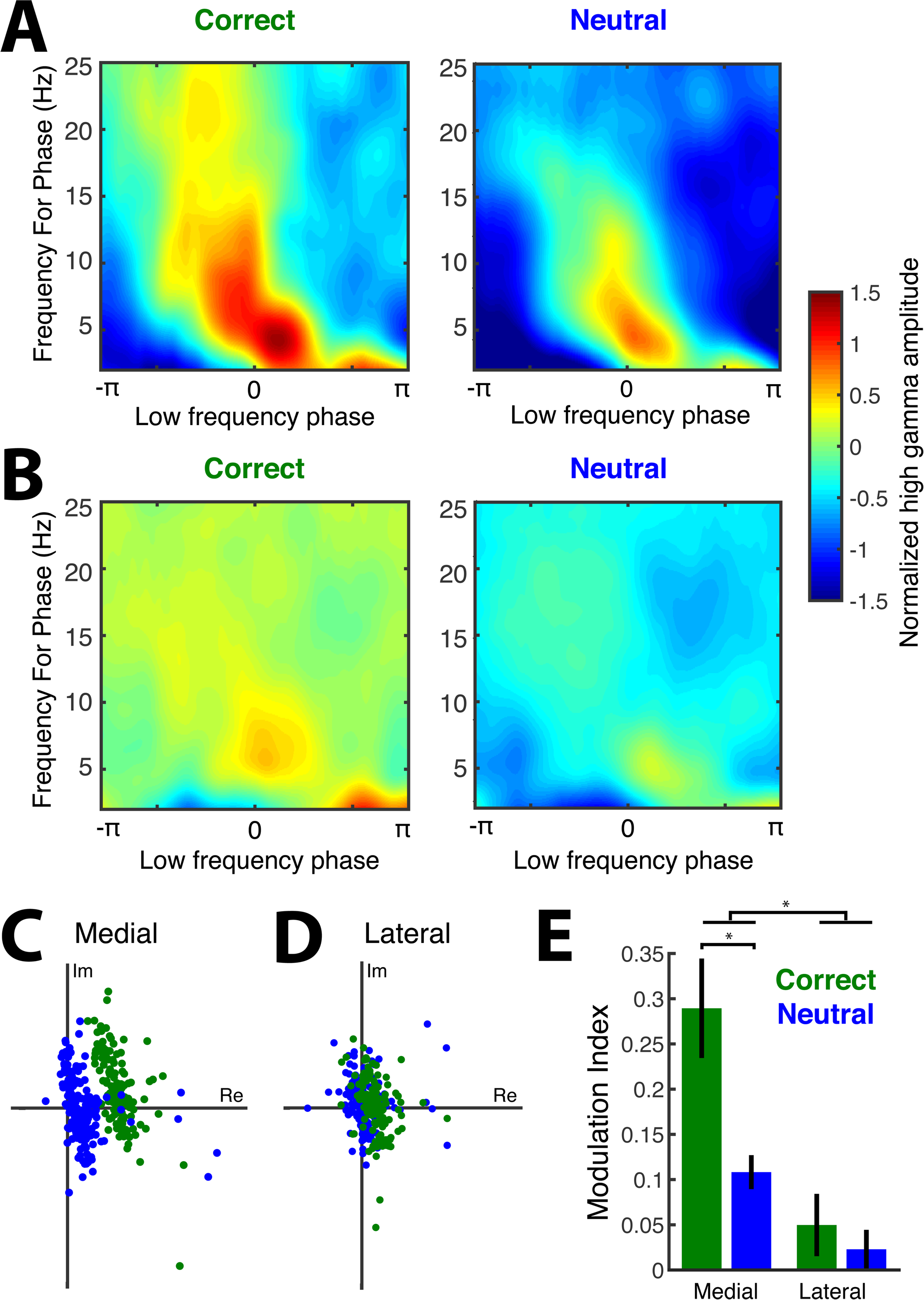
Phase-amplitude coupling (PAC) of low and high frequency LFPs. **A, B,** Representative PAC for a medial (**A**) and lateral (**B**) contact in one subject for each feedback condition (correct and neutral). **C, D,** Plots of the real (x axis) and imaginary (y axis) components of the modulation indices over trials for a representative patient for medial (**C**) and lateral (**D**) contacts. Blue dots represent the modulation indices for neutral trials and green dots represent the modulation indices for correct trials. **E,** Mean modulation indices across patients for medial and lateral contacts. Green and blue bars represent correct and neutral feedback trials, respectively. Asterisks indicate significant differences (p<0.01). Modulation indices were greater for correct feedback trials than neutral feedback trials in medial (Tukey-Kramer, p = 0.0052, N = 36 contacts), but not in lateral (Tukey-Kramer, p = 0.0598, N = 36 contacts) contacts. The difference in theta-high gamma coupling between correct and neutral trials was greater in medial contacts than lateral contacts (Tukey-Kramer, p = 0.0098, N = 36 contacts).

### Information transfer between dACC and dlPFC

The above results provide support for the hypothesis that the FRN is generated in the dACC, and that modulation of theta-gamma coupling in the dACC encodes feedback-related information. We next sought to determine the functional relationship between feedback signals across dACC and dlPFC, hypothesizing that these signals propagate from dACC to dlPFC.

To do so, we first used Granger causality (GC) techniques to study temporal correlations between dACC and dlPFC activty. To account for the non-stationarity of ECoG data, we compressed response-aligned broadband signals on all electrodes. Both medial-to-lateral and lateral-to-medial interactions were significant (*χ*^2^-Test, both p < 0.01, df = 6, corrected for 12 hypotheses using the FDR), although GC was insignificantly greater for medial-to-lateral interactions (Figure 5A). Peak spectral pairwise-conditional causalities were greatest in the low theta range (mean±std = 4.44±0.67 Hz) (Figure 5B). These results suggest that there are reciprocal interactions between dACC and dlPFC.

**Figure 5.**
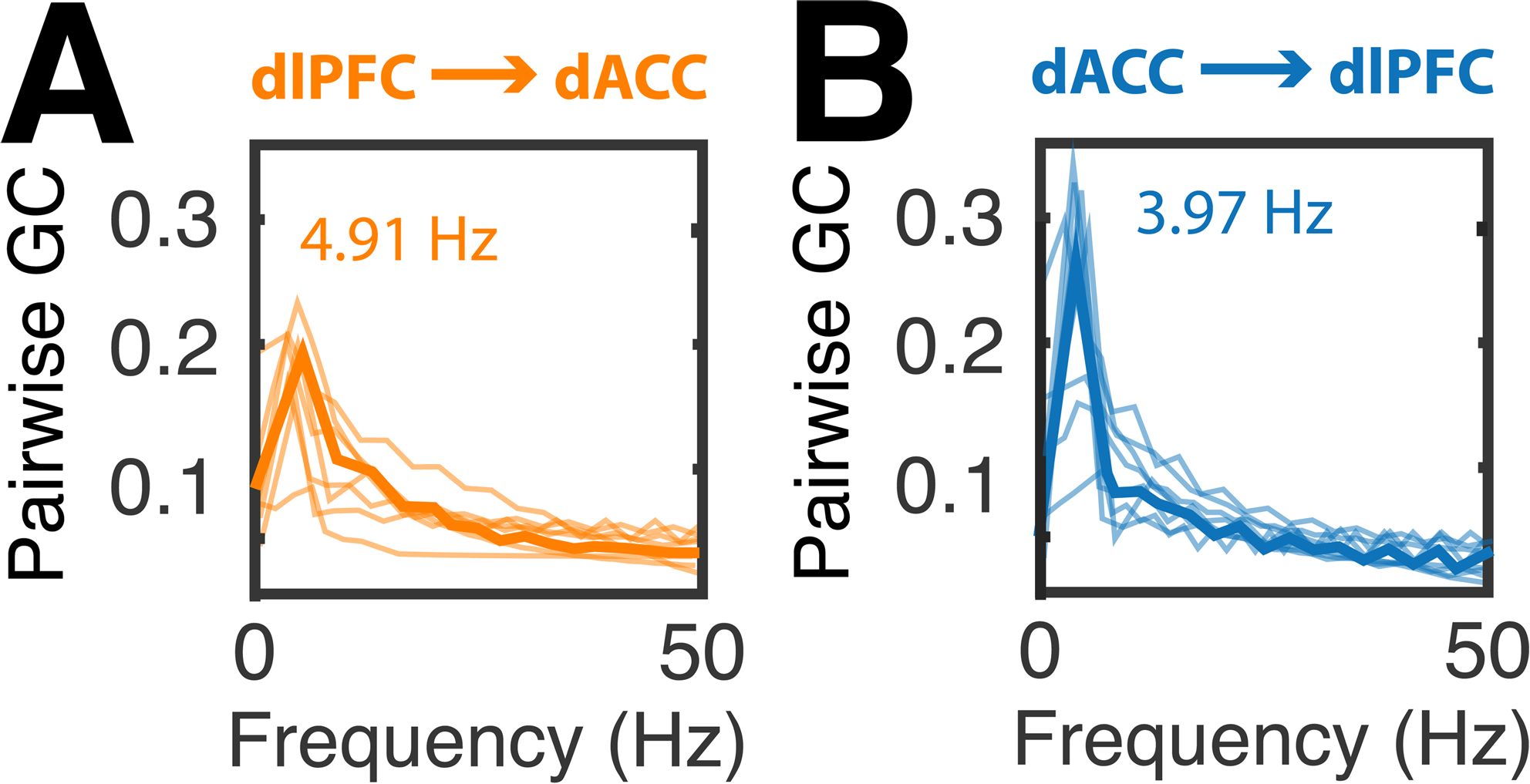
Pairwise conditional Granger Causality between dACC and dlPFC. **A,** Pairwise GC over frequencies for the lateral-to-medial direction. Mean GC values across patients are shown in dark bold orange, and patient specific data are shown in light orange. Peak frequencies averaged over patients are indicated in the colored text in the plot. **B,** Pairwise GC over frequencies for the medial-to-lateral direction. Mean GC values across patients are shown in dark bold blue, and patient specific data are shown in light blue. Peak frequencies averaged over patients are indicated in the colored text in the plot.

To avoid contending with the violations of GC assumptions typical of ECoG data, we further employed an analysis insensitive to these assumptions. We computed information transfer between medial and lateral PFC using a conditional mutual information (CMI) approach. CMI is a single-trial measure of the reduction in uncertainty about the variable of interest obtained by observing the current trial’s neural data from one contact given that one knows the response on another contact. In our case, CMI is a relative measure of the dependence of feedback valence representation in one cortical area on the other cortical area. Lateral-to-medial information transfer for each subject was not significantly greater than shuffled data after feedback presentation (*χ*^2^ test, all p > 0.01) and did not oscillate above a constant model (F-test, F = 0.288, p = 0.83) (Figure 6). However, CMI was significantly greater than shuffled data for medial-to-lateral information transfer for each subject (*χ*^2^ test, all p < 0.01), and oscillated (F-test, F = 36.50, p << 0.01) in the theta range (mean±std = 5.1 ±2.4 Hz) (Figure 6A,B). These results support the notion of a largely unidirectional dACC-to-dlPFC transfer of feedback-related information that oscillates with theta-range periodicity.

**Figure 6.**
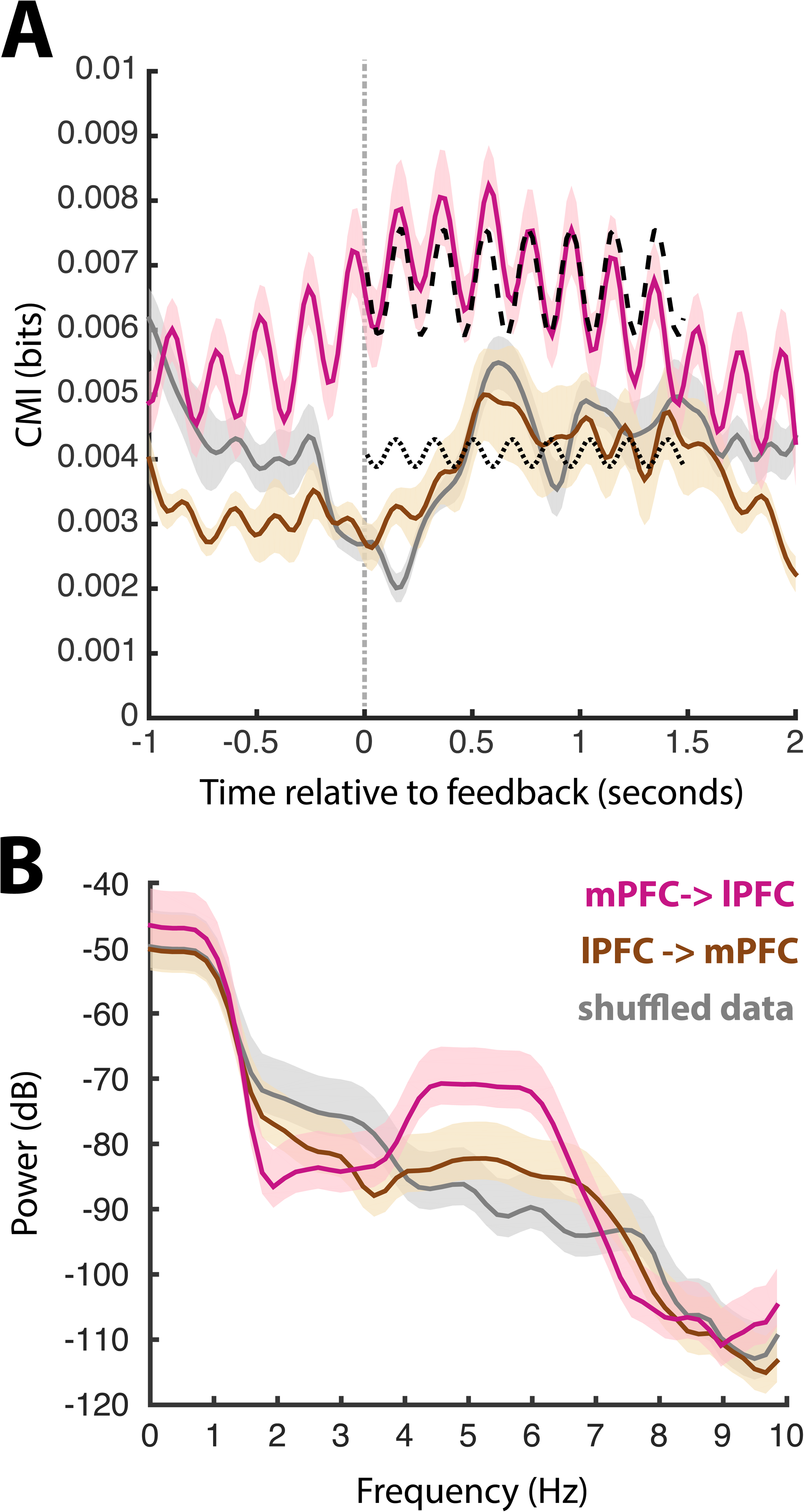
Conditional mutual information exchange between dACC and dlPFC. **A,** Mean CMI across subjects after feedback. Pink lines represent medial-to-lateral information transfer, and brown lines represent lateral-to-medial transfer. Gray lines represent shuffled data. Dashed and dotted lines show sinusoidal fits for medial-to-lateral and lateral-to-medial CMI, respectively. Vertical line represents feedback onset. **B,** Mean spectra of the CMI time series show increased theta power for medial-to-lateral CMI.

## Discussion

We examined the neural representation of prefrontal feedback signals using human ECoG with simultaneous medial and lateral PFC recordings. We report four main findings: 1) Low and high frequency ECoG signals from human PFC are sensitive to feedback valence. 2) These signals are larger and arise earlier in dACC compared to dlPFC. 3) Spectral analysis demonstrated that theta-gamma coupling modulation underlies these signals, and is also greater in dACC. 4) Information transfer analyses showed that feedback-related information is transferred from dACC to dlPFC with theta periodicity. These data thus provide direct evidence that the dACC is the cortical source of the FRN, and that theta modulation serves as a mechanism for communication of feedback-related information between dACC and lateral PFC.

### FRN localization: Direct support for a dACC source

Previous EEG source localization studies have posited various sites of origin for the FRN. Whereas the dACC is a common finding, others include the posterior cingulate cortex and basal ganglia (reviewed in (Walsh and Anderson, 2012)) All these structures receive input from midbrain dopaminergic cells, making them candidate regions for producing RPE signals. Source modeling of EEG has been helpful, but the requirement of solving the inverse problem from peripherally (scalp) recorded data makes identification of a single source difficult.

Our direct human intracranial recordings demonstrated the FRN at the level of individual electrode contacts in both low and high frequency ranges. Although signals were seen in both dACC and lateral PFC, those from dACC were larger and had shorter latencies. Human ECoG recordings have their own limitations: most importantly, that electrodes can only be placed based on clinical grounds. Because the basal ganglia are not a typical onset site for seizures, they are seldom placed there for epilepsy investigations. On the other hand, electrodes are frequently placed into the subthalamic nucleus and globus pallidus internus during deep brain stimulation procedures to treat movement disorders such as Parkinson disease and dystonia (Zavala et al., 2013; 2014). Investigation of feedback responses in these nuclei in humans is therefore possible, and will be the subject of future work.

A recent combined EEG-fMRI study modeled the fMRI response informed by the EEG and also identified the dACC as the FRN source (Hauser et al., 2013). Their dynamic causal modeling analysis also showed that FRN signals were not arriving to dACC from other structures. Thus taking the current study and previous studies together, there seems little doubt that the dACC is the source of the FRN.

### FRN dynamics: theta modulation

There is increasing evidence that the FRN and similar potentials, such as the conflict-elicited N2 and the error-related negativity, share a physiological basis, namely modulation of theta power (Holroyd et al., 2004a; Cavanagh and Frank, 2014). This commonality has lead some to propose the existence of an interrelated group of cognitive control signals known as frontal midline theta (FMΘ). FMΘ is an increase in theta-range (4-7 Hz) spectral power observed in the event-related potentials recorded on frontocentral EEG contacts. This group of FMΘ signals is thought to employ theta modulation as a means of communicating a certain class of information with other brain regions. FMΘ may be a signature of dACC activation in response to threats, either corporeal or cognitive, in order to signal the need for behaviorally relevant attentional control (Cavanagh and Shackman, 2014).

The current data show that the amplitude of high gamma activity, thought to be an indicator of local neuronal processing (Miller et al., 2009; Buzsáki et al., 2012), coupled to the phase of low frequency oscillations specifically in the theta band. The modulation of this phase-amplitude coupling was sensitive to feedback valence, and was stronger in dACC than in dlPFC. Furthermore, our information transfer analyses both showed peaks in the theta range. The linear, parametric analysis, GC, indicated reciprocal interaction between dACC and dlPFC that was dominated by the theta range. In the context of the current study, GC has some caveats. First, that we applied a compressor to the signal in order to achieve stationarity and second, that the duration of the responses, especially those high frequency responses lasting longer than 500 milliseconds, could bias the analyses. Conditional mutual information analysis, which is nonlinear and nonparametric, showed that feedback information arising in dACC propagated to dlPFC, and information from dlPFC did not propagate to dACC. Despite the inclusion of broadband LFP in this these analyses, the spectral composition of this information transfer showed a prominent peak centered at 5 Hz. All of these results support the contention that the dACC utilizes theta range oscillations to communicate with other cognitive control centers.

Our data thus lend support to the importance of FMΘ signals in general, and to the association of the FRN with this group in particular. The conception that FRN, and FMΘ, originates in the dACC, yet is represented in dlPFC is also consistent with our results. While some EEG studies have suggested that FMΘ arises through reciprocal activation of medial and lateral prefrontal cortex (Asada et al., 1999), the intracranial electrophysiology we present indicates a largely unidirectional, medial-to-lateral transfer of feedback-related information. Another EEG study suggested that FMΘ is associated with increased theta range synchrony among increasingly lateral scalp electrodes (Cavanagh et al., 2010). However, theta power on these lateral scalp electrodes correlated with neither prediction error nor behavioral adaptation (Cavanagh et al., 2010). Our results indicating that dACC sends information to lateral prefrontal areas via a theta rhythm could explain these EEG results.

Theta oscillations have also been observed in the dACC in other species. Simultaneous single neuron and LFP recordings have enabled investigation into the mechanism and function underlying theta modulation. Previous rat (Narayanan et al., 2013) and monkey (Womelsdorf et al., 2010) studies have demonstrated that medial PFC neuronal spiking synchronizes with theta rhythms, and the magnitude of this coherence correlates with behavioral measures of cognitive control. These results argue for a mechanism by which theta-band oscillations sharpen spiking output to particular phase intervals within a theta cycle (Womelsdorf et al., 2010). This mechanism of temporal focusing may synchronize distant regions within a cognitive control network and promote more efficient spike-based communication between them.

Spike-phase coupling has been previously described as a means of integrating brain-wide networks (Fries, 2005; Lakatos, 2005; Lakatos et al., 2009; Womelsdorf et al., 2010). One salient example of low-frequency LFP phase coordinating neuronal population activity in distributed networks showed that prefrontal neurons are sensitive to distinct, yet diverse array of rhythms (Canolty et al., 2010). As in the current study, theta band modulation has frequently been observed in PFC (Canolty et al., 2006; Voytek et al., 2010; Miller et al., 2012; Zavala et al., 2014). Whether the dACC establishes this rhythm, or it is entrained by another structure remains to be determined.

Theta rhythms are not unique to the dACC and to mechanisms of learning and cognitive control. It is known in humans and rodents that action potentials commonly couple to hippocampal theta rhythms, which are 4-8 Hz in the rat and 1-4 Hz in humans (Jacobs, 2014), and that this coupling likely supports memory processes (Buzsáki, 2005). Interestingly, action potentials couple preferentially to 4-6 Hz rhythms in the dACC in humans (Jacobs et al., 2007), which is the same range in which functional differences in phase amplitude coupling were observed in the current study. The medial-to-lateral CMI presented here also took on rhythmicity in precisely the 4-6 Hz range. These results support FMΘ arising from an information transfer architecture, as has been proposed in the rodent hippocampus (Mizuseki et al., 2009).

## Conclusions

We have shown that feedback is associated with broadband ERPs and high gamma responses in the dACC and dlPFC. We have also shown that these signals are coupled to theta-band oscillations, and that information is conveyed from dACC to dlPFC entrained to a theta rhythm. Our data support a novel model for the generation of FMΘ associated with feedback in human subjects, and pave the way for future experiments to test how humans use FMΘ signals to coordinate behavioral responses to feedback or other indications for the need for cognitive control. Together these results suggest that feedback-related information is initially processed in the dACC and quickly communicated to lateral prefrontal cortex as part of a cognitive control signal (Sheth et al., 2012; Shenhav et al., 2013). Lateral PFC presumably incorporates this information to influence ongoing behavioral modifications relevant to the immediate context and long-term goals (Miller and Cohen, 2001; Carter and van Veen, 2007).

## Acknowledgements

This work supported by NIH K12 NS080223 and the Dana Foundation. Thanks to Jennifer Russo for help collecting data. Thanks to Lucia Melloni, Saskia Haegens, Guillermo Horga, Nima Mesgarani, and Charles Schroeder for helpful developmental advice.

